# The structural basis for RNA selectivity by the IMP family of RNA binding proteins

**DOI:** 10.1101/618587

**Authors:** Jeetayu Biswas, Vivek L. Patel, Varun Bhaskar, Jeffrey A. Chao, Robert H. Singer, Carolina Eliscovich

**Affiliations:** Department of Anatomy and Structural Biology, Albert Einstein College of Medicine, Bronx, NY, 10461, USA; Department of Radiation Oncology, Massachusetts General Hospital, Boston, MA, 02114, USA; Friedrich Miescher Institute for Biomedical Research, CH-4058 Basel, Switzerland; Howard Hughes Medical Institute, Janelia Research Campus, Ashburn, VA 20147, USA; Department of Medicine, Albert Einstein College of Medicine, Bronx, NY, 10461, USA

**Keywords:** IMP2, ZBP1, RNA-binding protein, KH domain, variable loop

## Abstract

The Igf2 mRNA binding proteins (ZBP1/IMP1, IMP2, IMP3) are highly conserved post-transcriptional regulators of RNA stability, localization and translation. They play important roles in cell migration, neural development, metabolism and cancer cell survival. The knockout phenotypes of individual IMP proteins suggest that each family member regulates a unique pool of RNAs, yet evidence and an underlying mechanism for this is lacking. Here, we combine SELEX and NMR spectroscopy to demonstrate that the major RNA binding domains of the two most distantly related IMPs (ZBP1 and IMP2) bind to different consensus sequences and regulate targets consistent with their knockout phenotypes and roles in disease. We find that the targeting specificity of each IMP is determined by few amino acids in their variable loops. As variable loops often differ amongst KH domain paralogs, we hypothesize that this is a general mechanism for evolving specificity and regulation of the transcriptome.

## Introduction

Post-transcriptional regulation of RNAs is accomplished through their interactions with RNA binding proteins (RBPs). With over 1000 RBPs expressed in humans^1^, their ability to regulate RNA is extensive. RBPs participate in all aspects of an RNA’s life^1^, many are deposited as soon as the RNA is transcribed and they contribute to control over export, localization^2^, translation^3^ and decay^4^. Progress has been made in studying RBPs on the atomic level^5^, with single cell resolution^1^, as well as on a genome wide level by profiling the RNAs that interact with individual RBPs^6^. Many families of RBPs are involved in the regulation of normal and pathological processes, but how this regulation occurs or how differences in regulation amongst family members have evolved is not appreciated. Some prominent RBP families that have been studied on both a structural and genome wide level include Fragile X proteins^7–9^, NOVAs^10–12^ and IGF2BPs^1,13–15^.

The IGF2BP family (IMP) consists of highly conserved RBPs^1^. The founding member of the IMP family, ZBP1, was first characterized in chicken embryonic fibroblasts and then subsequently its paralogs were discovered and named CRD-BP, IMP1, IGF2BP1 and VICKZ1. While the naming convention varied according to the lab that discovered the protein, the functions of ZBP1 as well as amino acid sequence identity are highly conserved across species. In mammals, there are three IMP family members (ZBP1, IMP2 and IMP3). Each protein contains six canonical RNA binding domains, two RNA recognition motif (RRM) domains and four hnRNP K homology (KH) domains. These RNA binding domains are arranged into three pairs (RRM12, KH12 and KH34). In humans, IMP members share an overall sequence identity of 56%. The percentage increases to 70% when unstructured linker regions are excluded and the comparison is limited to the individual RNA binding domains^16^. Though these proteins share similar amino acid sequences, each IMP family member regulates a different pool of cellular RNAs and knockouts (KO) of ZBP1, IMP2 and IMP3 have identifiable phenotypes^17,18^.

It has been shown that ZBP1 binds to the 3’ UTR of ß-actin mRNA^19^, and in so doing, prevents its translation until it reaches its destination^1^, be it the leading edge of a fibroblast^1,20,21^ or a synaptic spine destined for remodeling^19,20,22^. ZBP1KO mice are perinatal lethal, with gross developmental abnormalities, consistent with the observation that ZBP1 expression is maximal during the period of mid to late embryonic development^18^. To identify targets of ZBP1, Systematic Evolution of Ligands by Exponential Enrichment (SELEX) followed by biochemical and structural characterization revealed two RNA elements^13,23^. The identity of RNA bases and the spacing between them was critical for recognition ^1,24^. Genome-wide searches using these stringent binding criteria identified mRNA targets that are critical for cell growth, organization and neural development, data that is consistent with the phenotype of the ZBP1KO mouse^13,18^. These studies provided a list of targets that are critical for ZBP1’s roles in fibroblast migration^25^, cancer metastasis^26,27^ and synaptic plasticity^1,1,22^.

Investigation of IMP2 has been less extensive than for ZBP1 and IMP3. ZBP1 and IMP3 share high sequence identity (73% overall) and common temporal expression patterns (mid to late period of embryonic development, with little to no expression in the adult). Clinical studies have found their expression to be reactivated in a range of tumors, hinting at a potential role for ZBP1 and IMP3 in cancer pathogenesis^16^. IMP2 shares less sequence homology with the other two family members and its expression persists throughout life^28^. Recently, IMP2KO mice have been observed to have a metabolic phenotype that extends their lifespan and renders them resistant to high fat diet induced obesity and type-II diabetes^17^. Further corroborating the role for IMP2 in metabolism, studies have shown that IMP2 regulates the translation of subunits critical for oxidative phosphorylation in normal cells as well as in glioblastoma^29^. Moreover, both IMP2 and a splice variant (p62) appear to be harbingers of a poor prognosis in gastrointestinal, hepatocellular and breast carcinomas^30–32^, possibly through its regulation of mitochondrial function^17,29^.

Though much work has gone into studying individual family members of the IMP family, it is not clear if they recognize similar or different targets. While genome wide CLIP studies show a correlation between the total RNA pools targeted by each protein^14,15^, we investigated individual targets and find, while some targets can bind to both ZBP1 and IMP2, others are specific for one or the other. The similarities and differences between the binding elements of ZBP1 and IMP2 reveal how RNAse digestion based techniques (such as CLIP) can digest away a necessary “GG” motif that is common to the consensus RNA binding sequence of both proteins. Here, utilizing both crystallography and NMR spectroscopy revealed how divergence in amino acid sequence can contribute to differences in RNA specificity. Targeted mutations in the amino acids of IMP2 allow the mutant to bind to ZBP1 targets. This represents how engineering of KH domains can provide insights into how this highly conserved family of RNA binding proteins has evolved to express different genes post-transcriptionally within the cell.

## Results

### Structural similarity of ZBP1 and IMP2 KH34 domains

ZBP1 and IMP2 have high sequence identity (>70%) in their RNA-binding domains (Fig. 1a). *In vivo* functional studies as well as *in vitro* work suggested that KH34 domains are responsible for RNA recognition^1,24^. To determine if there are differences in the overall topology of the KH34 domains, we first crystallized the KH34 domain of IMP2 and compared it to the previously determined structure of ZBP1^1^ (Fig. 1b).

**Fig. 1.**
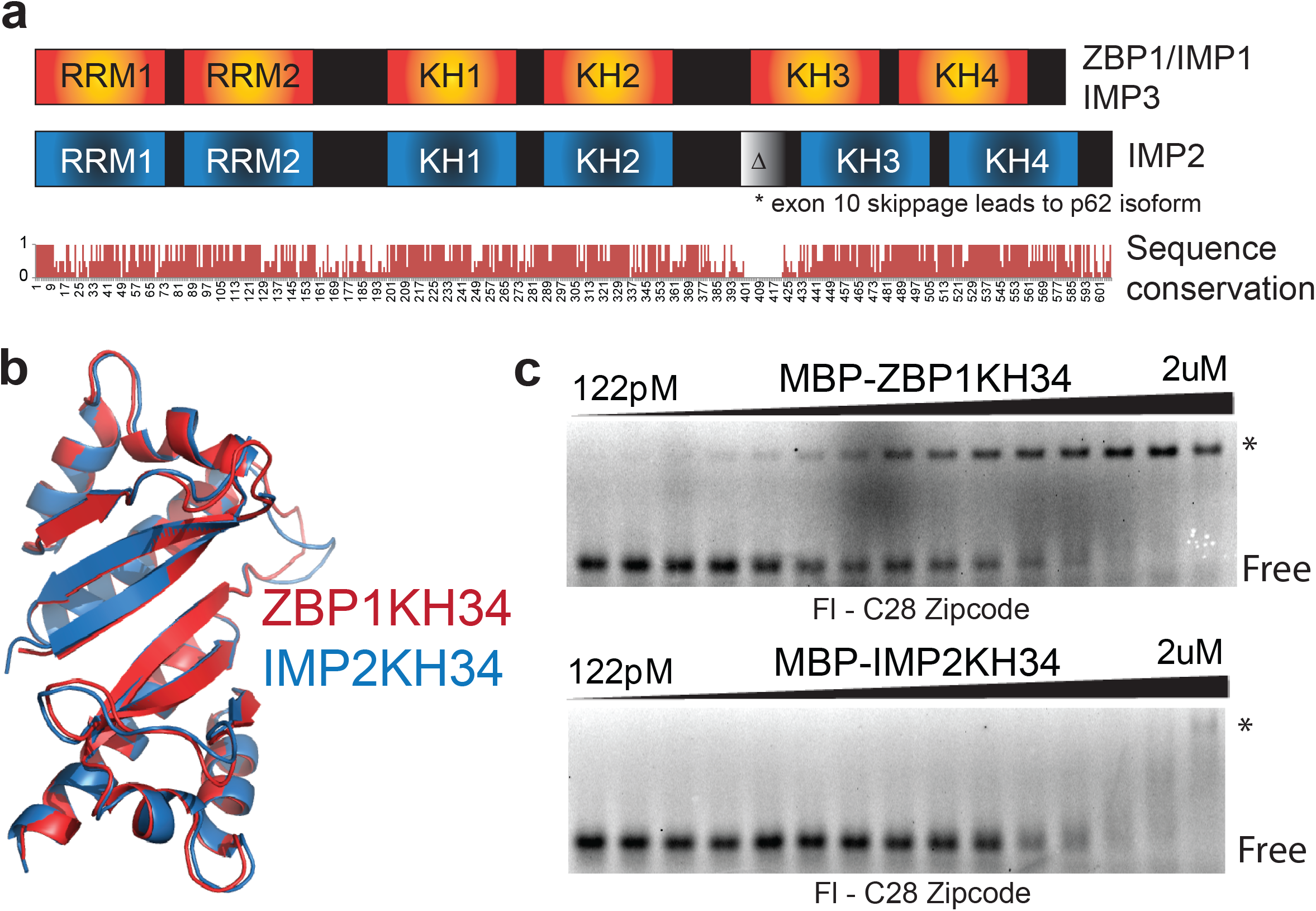
ZBP1 and IMP2 overview and crystal structures. **a.** Overall domain structure and sequence conservation of the IMP family members above. Below, sequence conservation across the IMP family members. **b.** Ribbon diagrams overlaying the crystal structures of ZBP1KH34 in red (PDB: 3KRM) and IMP2KH34 in blue (this study). **c.** Representative EMSAs for wild-type C28 ß-actin zipcode RNA. The filled triangle represents a 1:1 serial dilution of ZBP1KH34 (top) and IMP2KH34 (bottom). The RBP-RNA complex (*) and free RNA (FREE) are labeled.

Similar to ZBP1KH34, we found that the individual KH3 and KH4 domains of IMP2 adopted the type 1 KH fold (βααββα) and were also arranged in an anti-parallel pseudo-dimer orientation. The overall structures of IMP2KH34 and ZBP1KH34 are almost identical (RMSD of 0.5 Å for all C_α_ atoms) (Fig. 1b). Because the anti-parallel pseudo-dimer of ZBP1KH34 recognizes a bipartite RNA element^1^, we hypothesized that IMP2KH34 may recognize a similar bipartite RNA topology. However, it is not known if the structural similarity of IMP2KH34 allows it to recognize the same RNA-binding element as ZBP1.

### IMP2 does not bind to the canonical RNA target of ZBP1

To determine if IMP2KH34 could recognize the canonical RNA target of ZBP1KH34, we tested its binding affinity using quantitative Electrophoretic Gel Shift Assay (EMSA). We used the minimal ß-actin zipcode element as an *in vitro* RNA (C28 zipcode RNA^1,13^). We observed that IMP2KH34 had lower affinity to the ß-actin C28 zipcode RNA (Kd = 161nM) (Fig 1c, Sup Fig. 5) compared with ZBP1KH34 (Kd = 15nM) (Fig. 1c). This result suggested that the KH34 domains of IMP2 and ZBP1 might recognize and bind different RNA targets.

### SELEX identifies IMP2 specific RNAs

To understand if the inability of IMP2KH34 to bind the ß-actin zipcode RNA was due to a specific sequence preference or a general RNA binding defect in the KH34 domains of IMP2, we performed SELEX using IMP2KH34 as a protein bait. We generated an RNA library where 30 nucleotides were randomized at each position (N30). After *in vitro* transcription, the N30 pool was passed over an amylose resin bound to a recombinant MBP-IMP2KH34 fusion protein. After 9 rounds of selection, we found that the RNA library was bound to IMP2KH34 with an approximately 800-fold higher affinity compared with the initial randomized RNA library (Fig. 2a, b). From the last round of selection, we identified a bipartite RNA element that included a 5’ CA element, CU<u>CA</u>C, followed by ten nucleotides and then a variable 3’ GG element, (A/U)-<u>GG</u>-(A/U). These CA and GG recognition elements exhibited similarity to the ZBP1KH34 consensus, C<u>GG</u>AC and (C/A)-<u>CA</u>-(C/U)^1,13^ (Fig. 2a). This indicates that the SELEX procedure identified closely related sequences from two different members of the IMP family.

**Fig. 2.**
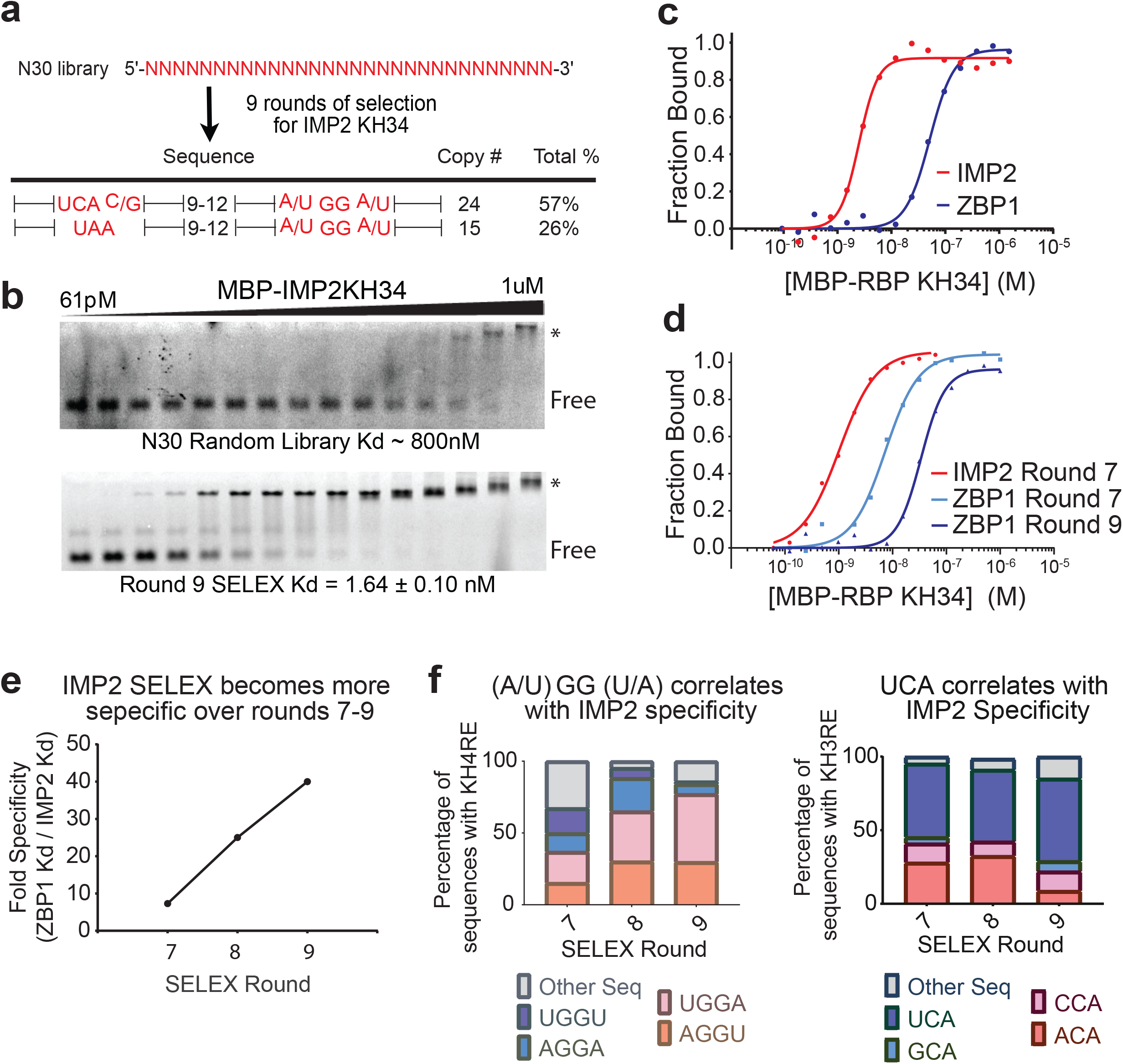
SELEX discovers targets that are IMP2 specific. **a.** The sequences of RNAs after 9 rounds of SELEX are shown. The range of nucleotide spacing between the nonrandomized IMP2 recognition elements is indicated for each sequence. Copy number and percentage of pool are listed. **b.** Representative EMSAs for N30 SELEX library (top) and round 9 SELEX library (bottom). The filled triangle represents a 1:1 serial dilution of IMP2KH34. The RBP-RNA complex (*) and free RNA (FREE) are labeled. **c.** Quantification and fit to the Hill equation of representative EMSA results for IMP2KH34 (solid red line) and ZBP1KH34 (solid blue line) binding to the round 9 SELEX library pool. **d.** Quantification and fit to the Hill equation of representative EMSA results for IMP2KH34 (Round 7, solid red line) and ZBP1KH34 (round 7, solid cyan line, round 9, solid blue line) binding to different SELEX library pools. **e.** Quantification of SELEX specificity for IMP2KH34 within each round SELEX library pool. The library specificity was calculated as the ratio between the Kd of ZBP1KH34 and the Kd of IMP2KH34 at a particular round of selection. **f.** After sequencing each round of SELEX, the individual “GG” (left) and “CA” (right) motif occurrences were counted as a percentage of total SELEX sequences. The four most abundant motifs were plotted in terms of their relative abundance in each of the sequenced SELEX rounds. Source data are provided as a source data file.

To demonstrate that the CA and GG elements found in the SELEX procedure were both necessary for binding to IMP2KH34, *in vitro* transcribed RNAs with either the CA or GG sequences mutated to CG or AA, respectively, were tested for binding by IMP2KH34. We observed that the mutation of either the CA or the GG element completely abolished binding of the RNA to IMP2KH34 (Sup Fig. 1a, b). These results confirmed that the 5’ CA and 3’ GG elements were required for IMP2KH34 specific binding.

We then tested representative sequences from the last round of selection. These RNAs showed consistently tight binding to IMP2KH34 and not to ZBP1KH34 (Sup Fig. 1c, d). These results suggested that the sequences conserved (the CA and GG elements and their spacing) across different RNAs mediated the IMP2KH34 specificity of the SELEX targets.

To determine the evolution of specificity, the affinity of individual RNA pools to IMP2KH34 at each SELEX round was tested. As a control, the same pools of RNA were tested against ZBP1KH34 (Fig. 2d, e). We observed that after four rounds of selection, IMP2KH34 bound with higher affinity to the RNAs (Kd = 2.4 nM) compared with ZBP1KH34 (Kd = 8 nM) (Sup Fig. 2a, b). The specificity of the library (defined as the ratio of the IMP2KH34 Kd / ZBP1KH34 Kd) increased across the subsequent rounds of selection (Fig. 2c, d and e). This increase in specificity was due to an increase in pool’s affinity for IMP2 and a decrease of the pools ability to bind ZBP1 (Fig. 2d). This suggests that during early rounds in SELEX, sequences can bind to both IMP2KH34 and ZBP1KH34 but become enriched for sequences that favor the binding of IMP2KH34 relative to ZBP1KH34 (Fig. 2c).

Both CA and GG dinucleotides are necessary for binding to all the IMPs^1,13,33^ and were enriched by SELEX in this study. KH domains recognize 4-6nt sequence elements^5^; the IMP KH domains have been shown to recognize 4nt sequences^1,13,33^, therefore the nucleotides flanking the CA and GG motifs are likely important for binding. Their selective enrichment also changed across the last three rounds of SELEX (i.e., rounds 7, 8 and 9) (Fig. 2e, f). We found that RNAs containing the 5’ UCA and 3’ (A/U)-GG-(A/U) sequence were enriched; AGGU and UGGA motifs had the highest enrichment (Fig. 2f). In contrast, the GG element present in the ß-actin zipcode RNA must be CGGA^13^ in order to bind ZBP1. Therefore, the enrichment of AGGU and UGGA elements as IMP2 targets may provide an explanation for IMP2KH34 binding specificity. Altogether these results suggest that IMP2KH34 may favor binding to AGGU and UGGA, which are disfavored by ZBP1KH34.

### NMR Spectroscopy maps RNA binding interface of IMP2KH34

To determine the amino acids of IMP2KH34 that interact with the RNA, ^15^N-HSQC spectra were collected of KH34 in complex with short RNA sequences containing either the 5’ RE CCC<u>UCA</u>CC or the 3’ RE UU<u>UGGA</u>AC (Fig. 3 and Sup Fig. 3a, b, e, f). We found that the 5’ CA and the 3’ GG RNA element were bound specifically by the KH3 and KH4 protein domain respectively (Fig. 3a and Sup Fig. 3c, d, e, f). The chemical shift perturbations of the amino acids involved were near the opposite ends of the protein, a majority of which were near the GXXG motif and the variable loop (Fig. 3b and Sup Fig. 3b, e, f). This result was consistent with ZBP1 KH domain/RNA interaction where the highly conserved GXXG motif faces the phosphate backbone and the variable loops face the nucleobases^13,34^. Together the two form a vice like structure that binds to the RNA^5^. Altogether these results confirm that NMR spectroscopy can identify the same binding region in all IMP family members.

**Fig. 3.**
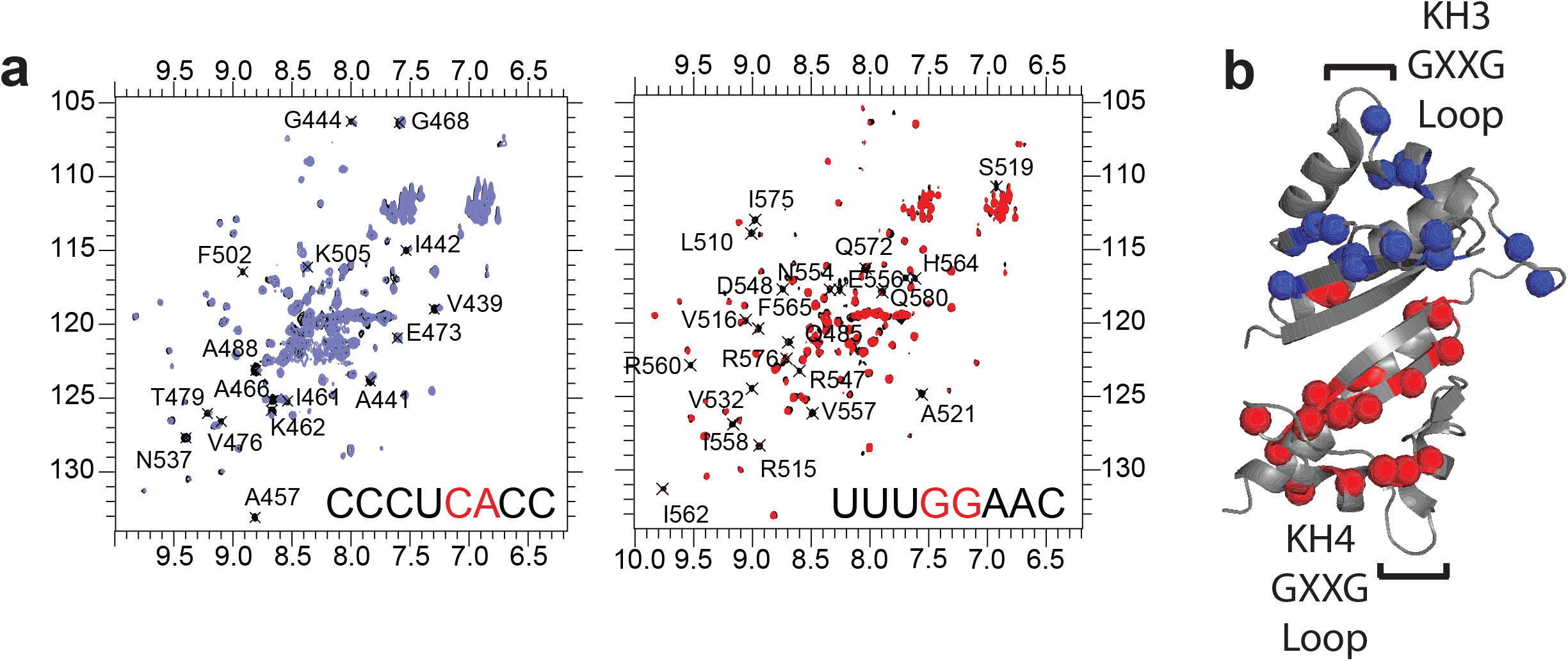
IMP2 recognizes specific RNAs through interactions between the GXXG motifs and the variable loops. **a.** ^1^H ^15^N HSQC spectra of IMP2KH34 showing residues perturbed (black text) during separate titrations of the CA (left, blue) and GG (right, red) element containing RNAs. Sequence of the RNA used is depicted on bottom right of each spectrum. Enlarged spectra of the titrations are shown in Sup Fig. 3e. **b.** Location of amide resonances altered upon binding to the CA element containing RNA (blue) and GG element containing RNA (red).

### IMP2KH34 variable loop mutagenesis mimics ZBP1 binding

Mutation of the GXXG motifs completely abolishes all nucleic acid binding^35^. We hypothesized that the highly conserved GXXG motifs mediate phosphate backbone interactions and that the variable loops functioned to recognize specific nucleobases. To determine if the variable loops mediate differences in RNA binding between ZBP1KH34 and IMP2KH34 we generated a chimeric protein where amino acids present near both variable loops of IMP2KH34 were exchanged with the corresponding amino acids present in the same position on ZBP1KH34 variable loops (Fig. 4a and Sup Fig. 4a). We found that this exchange converted IMP2 into a ZBP1 like binding protein, which bound to the ß-actin zipcode, the ZBP1 RNA binding element (Fig. 1c and Fig. 4b). This result indicates that the variable loops mediated the RNA binding specificity of IMP2KH34. To further determine if both KH3 and KH4 variable loops were necessary and sufficient to mediate specific binding to RNA, we generated single variable loop mutants of either IMP2KH3 or IMP2KH4 (Sup Fig. 4a). We observed that only four amino acid mutations in the KH3 domain were necessary to allow IMP2 to bind to the ß-actin zipcode with nearly equal affinity to ZBP1KH34 (Fig. 4b-d). These results provide direct evidence that the variable loops of a protein KH domain can specify RNA sequence preference.

**Fig. 4.**
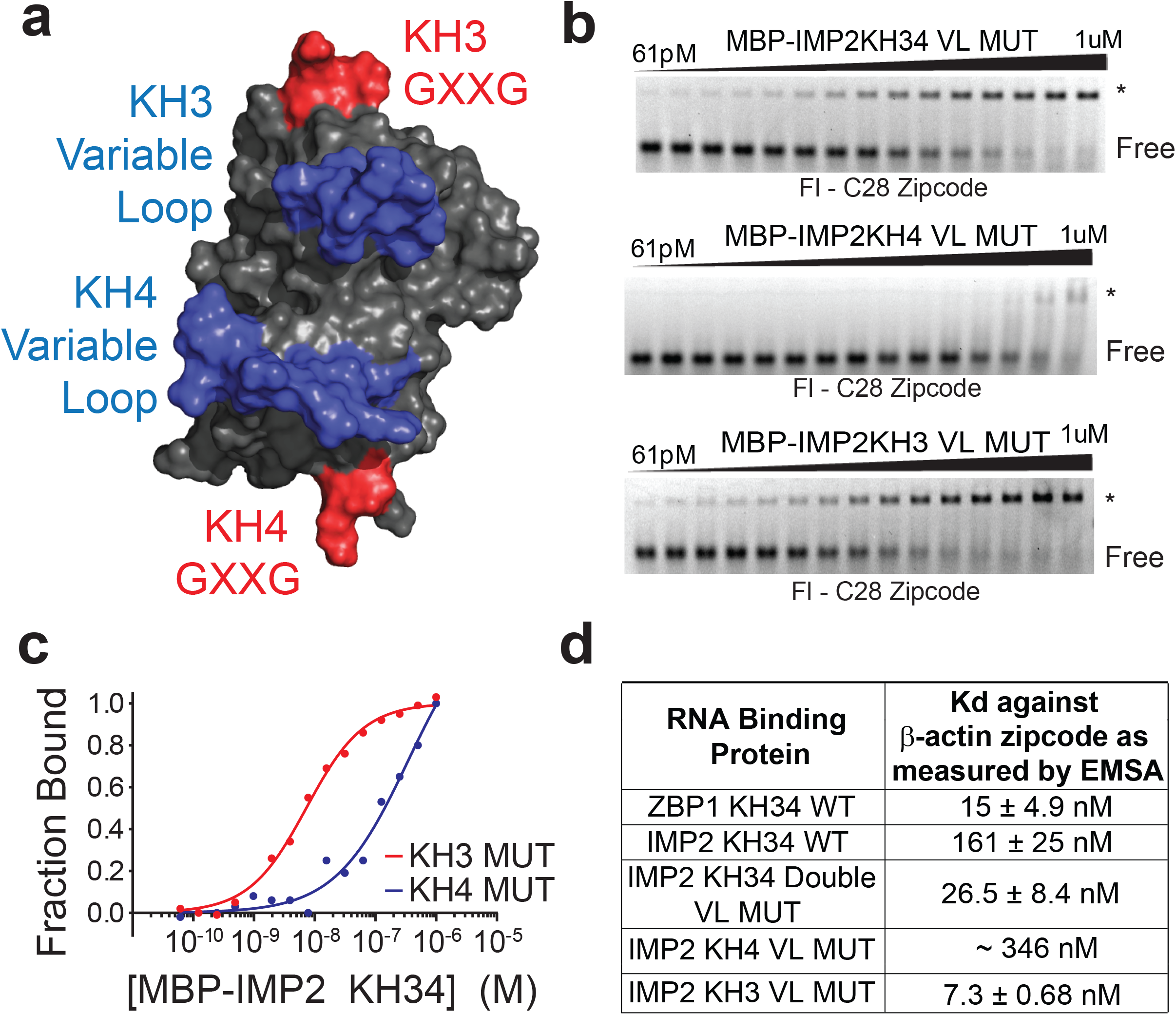
Mutations of the KH3 Variable loop are sufficient to determine RNA binding. **a.** Surface rendering of IMP2KH34 shows putative binding site of RNA with variable loops in blue and GXXG motifs in red. **b.** Top – double loop replacement allows for IMP2 to now bind to the ß-actin zipcode. Middle – The KH4 loop swap by itself does not increase affinity to the ß-actin zipcode compared to WT IMP2KH34. Bottom – KH3 Variable loop replacement is sufficient to swap the specificity of IMP2KH34 and allow it to bind to ß-actin zipcode. **c.** Quantification and fit to the Hill equation of top and middle representative EMSA results (in B) for IMP2KH3 VL MUT (solid red line) and IMP2KH4VL MUT (solid blue line) binding to ß-actin zipcode. **d.** Dissociation constants (Kd) of ZBP1KH34 WT (gel in Fig. 1c), IMP2KH34 WT (gel in Fig. 1c) and the IMP2 variable loop mutants (i.e., KH34 double VL MUT, KH3 VL MUT and KH4 VL MUT) and ß-actin zipcode measured by EMSA.

### IMP2 recognition element mutations reveal binding preference

The KH3 mutant was sufficient to confer a gain of RNA-binding function for IMP2 binding to the ß-actin zipcode. As evidenced by NMR chemical shift perturbations of IMP2 and ZBP1, both proteins’ KH3 domains interact with a CA motif. We showed that the CA dinucleotide core of the IMP2 5’ RE is also present in the ß-actin zipcode (Fig. 3a, b and Patel *et al*., 2012), therefore we determined if the two nucleotides adjacent to the CA in the ß-actin zipcode specifically inhibited IMP2 binding. We systematically mutated the individual nucleotides within and around the CA region of the ß-actin zipcode (Fig. 5a) and determined if the mutated ß-actin zipcode was bound to IMP2KH34. We found that mutagenesis of the ß-actin zipcode at positions adjacent to the CA motif, specifically position 22 A to U, increased the affinity of the ß-actin zipcode to IMP2KH34 by 10 fold (Fig. 5a-c). When nucleotides adjacent to the GG motif of the ß-actin zipcode were mutated binding for IMP2KH34 was not observed (Sup Fig. 5). Concordant with the SELEX results (Fig. 2a), the strongest binding occurred when position 22C was mutated to U, leading to a 5’ RE of UCA for IMP2KH34 (Fig. 5b, left, Fig. 5c). These results indicate that, unlike ZBP1, which can bind to both ‘(A/C)-CA’, IMP2 has a higher specificity for nucleotides in its 5’ RE, requiring the sequence UCA.

**Fig. 5.**
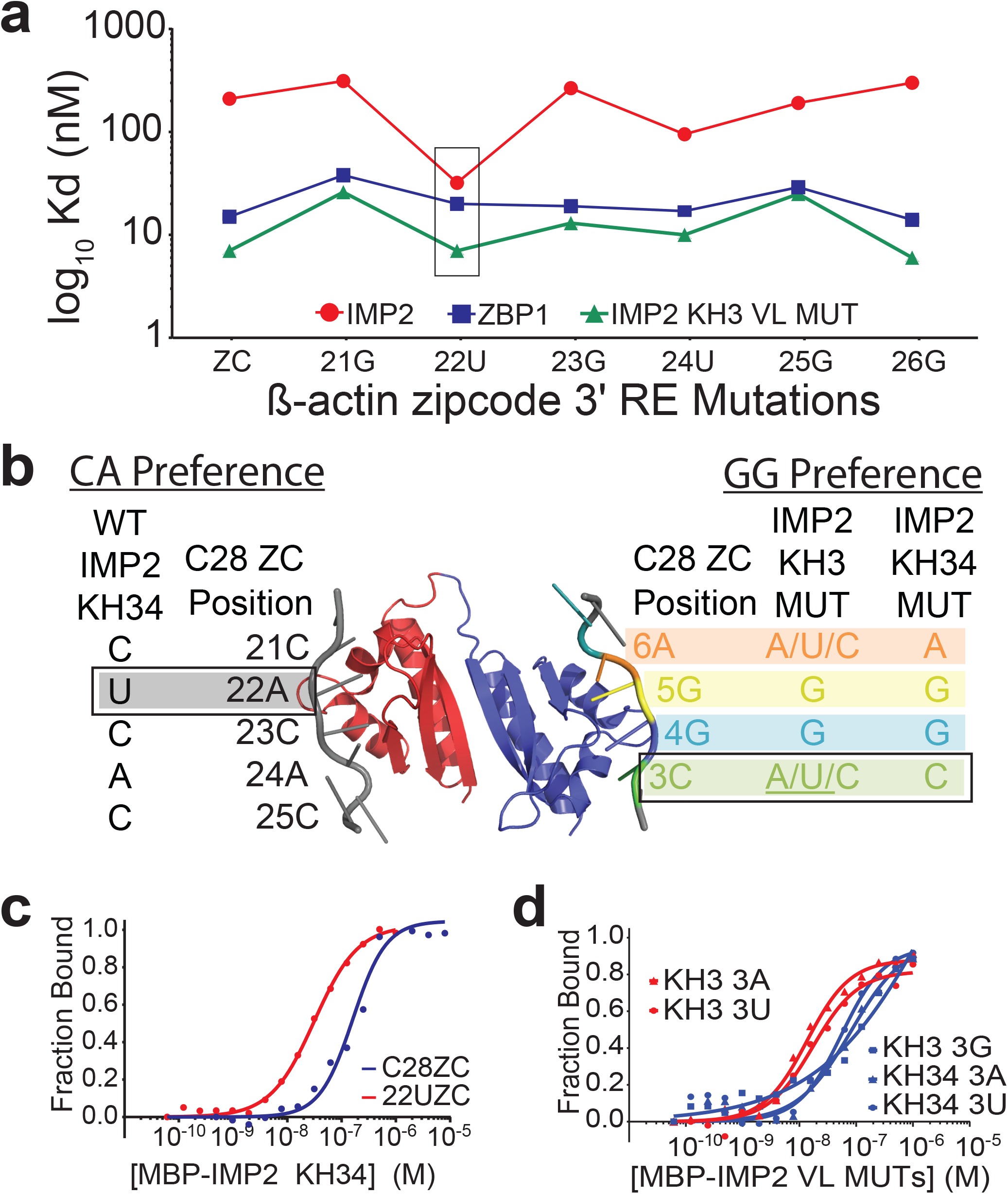
Mutations validate the sequence preference of individual IMP2 RNA recognition elements. **a.** Kd measurements of IMP2KH34 WT, ZBP1KH34 WT (from Patel et al., 2012) and IMP2KH3 variable loop mutant. Point mutations were made to the ß-actin actin zipcode sequence (X axis) and Kd was measured by EMSA (gels in Sup Fig. 5). **b.** Summary of sequence preferences for WT and variable loop mutants. Single nucleotide mutants of the ß-actin zipcode consensus elements were tested against IMP2 variable loop mutants and the preferred nucleotide at each position in written (gels in Sup Fig. 5). Data for ZBP1 consensus sequence preference was obtained from Patel et al., 2012. **c.** Quantification of representative EMSAs corresponding to B (grey box, gels in Sup Fig. 5). **d.** Quantification of representative EMSAs corresponding to B (green box, gels in Sup Fig. 5) showing differences between mutant zipcode sequences that bind (red) and those that do not bind (blue).

To determine the binding specificity of the KH4 domains, a similar approach measured binding of the KH3 mutant to the ß-actin zipcode (by now allowing for binding to CCA element) (Fig. 5b, right, Fig. 5d). The GG motif was crucial, and any mutation caused a complete loss of binding (Sup Fig. 5). The surrounding nucleotide preferences were different for ZBP1KH34 and the IMP2KH3MUT. ZBP1KH34 was intolerant to mutations surrounding the GG element, however the IMP2KH3MUT was permissive to almost any nucleotide except for G (Fig. 5d, Sup Fig. 5). To determine if the variable loops were sufficient to dictate this specificity we also repeated the experiment on the IMP2KH34 mutant and found it resembled ZBP1’s CGGA specificity. This suggests that ZBP1 has stringent sequence specificity for its KH4 domain whereas IMP2 is flexible and could tolerate any nucleotide except for a third G (Fig. 5b and Fig. 5d); and that the variable loops were sufficient to determine KH specificity.

### MP2 targets are highly enriched for metabolic functions

To identify potential mRNA ligands of IMP2KH34, we queried the 3’ UTRs of human and mouse transcripts for the bipartite IMP2KH34 RE defined in Fig. 2a. The lower and upper bound between these two elements was set to 10 and 15 nucleotides based on evidence from the structural constraints identified for ZBP1KH34^1^. We identified 2,790 human and 2,138 mouse 3’ UTRs containing the bipartite IMP2KH34 RE. To enrich for *bona fide* IMP2 mRNA targets, we used the overlap of these two lists to identify 503 genes that had the bipartite IMP2KH34 RE evolutionarily conserved (Fig. 6a).

**Fig. 6.**
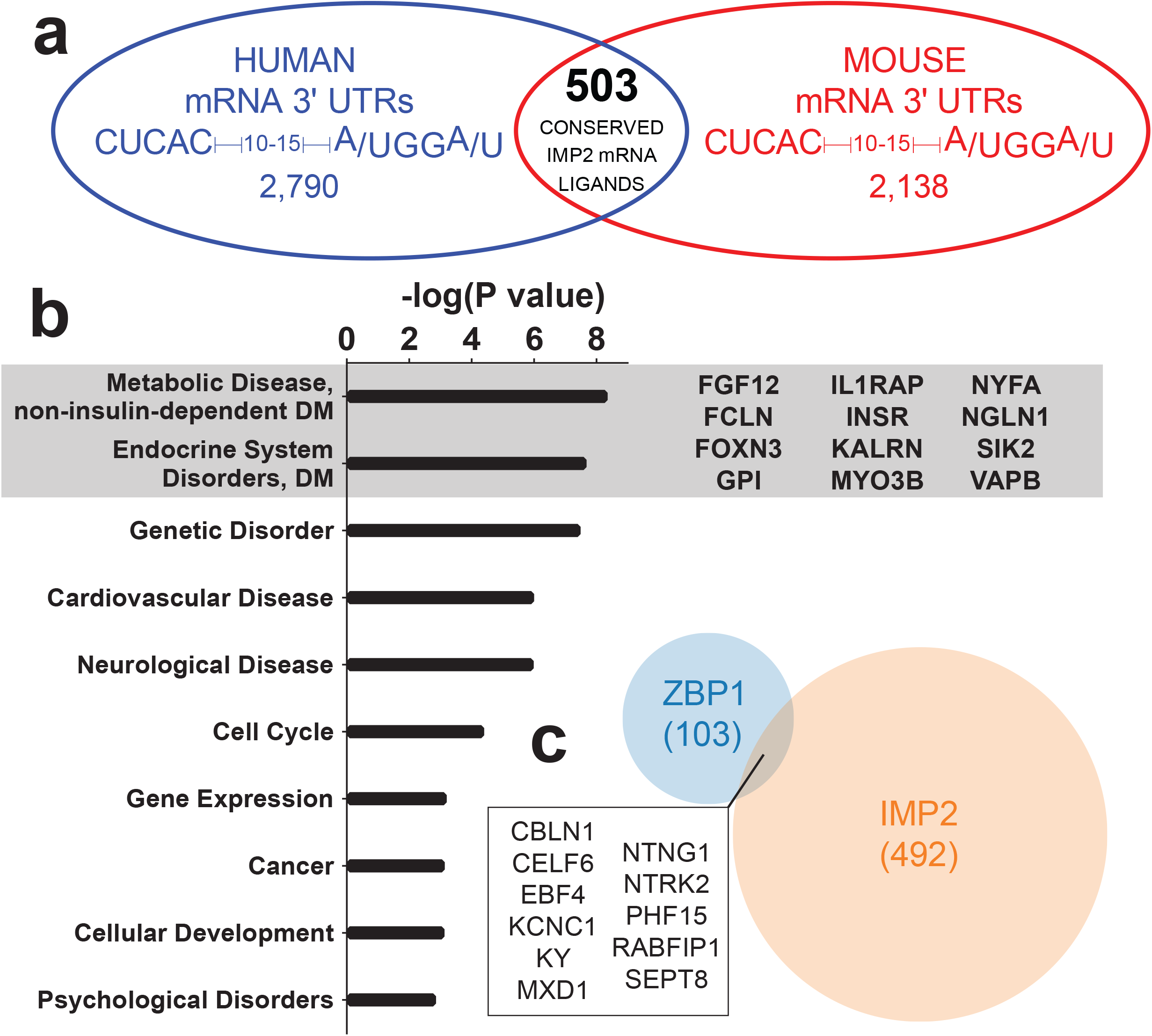
Genome wide search across 3’ UTRs for the IMP2 consensus sequence. **a.** Human and mouse mRNA 3’ UTRs containing the IMP2KH34 binding consensus sequences. Conserved RNA targets were used for gene ontology analysis in b. **b.** Gene ontology analysis of conserved mRNA ligands containing the bipartite IMP2KH34 RE. Grey box highlights gene symbols associated with diabetes. **c.** Conserved targets for ZBP1KH34 (Blue) and IMP2KH34 (Orange) show little overlap. Box highlights gene symbols that are predicted to bind to both ZBP1 and IMP2.

To define RNA targets further, we compared these transcripts to mRNA targets of ZBP1^13^. Pools of RNAs were ZBP1 or IMP2 specific, or potentially overlapping (Fig. 6c). In H9ES cells, RNAs bound by both ZBP1 and IMP2 (as discovered by eCLIP) provided evidence of transcripts such as SEPT8 being bound *in vivo* by both proteins^15^ (Sup Fig. 6). To show that our evolutionarily conserved IMP2 targets were conserved between mouse and human cells, published RIP seq data from both HEK293T^36^ and mouse brown adipose tissue^17^ were queried. In both experiments, the IMP2 consensus sequence containing RNAs show significant (P < 0.0001) enrichment over input or IgG controls (Sup Fig. 7a, b) and 189 RNA targets were enriched in both experimental samples (Sup Fig. 7c).

To associate these predicted RNA targets with functional annotations, we use gene ontology analysis to probe for the most significantly enriched functions and disease associations in the RNA ligands conserved between mouse and human. Gene ontology analysis of IMP2 target transcripts revealed an enrichment of genes associated with metabolic disease (Fig. 6b); enrichment was most significant for genes implicated in progression of type II diabetes (Supplementary Data 1). Therefore, the array of mRNA ligands potentially recognized by IMP2, such as the insulin receptor, may inform the mechanism of its association with type II diabetes in humans.

## Discussion

This work has shown that KH domains can confer sequence specific recognition of RNAs through their variable loop regions. We have shown that mutations within the variable loop regions are responsible for the divergence in consensus sequence between ZBP1 and IMP2. Mutating the variable loop amino acids as well as the nucleotides that they recognize has clarified differences in the sequence specificity between the most distant IMP family members.

Recently structural studies have investigated the binding preferences for the remaining family member, IMP3, and found an analogous bipartite consensus sequence of CGGC – 12 to 25 – CA^33^. Given that the amino acid sequence of IMP3 is closer to ZBP1 than IMP2, one may expect its consensus sequence to more closely resemble ZBP1’s CGG-(A/U) motif than IMP2’s (A/U)-GG-(A/U) motif^13^. This points towards a consistent evolutionary divergence amongst IMP family members for novel recognition sequences (Sup Fig. 8). Interestingly, in the study of IMP3, the authors found that KH12 required the same GG and CA consensus motifs as KH34. While other work has shown that KH34 are the major RNA binding domains for each of the IMP family members^1,24^, this brings up the possibility that their interactions with RNA targets may either be reinforced or modified by the presence of additional RNA binding domains. Modification of binding preferences by additional domains may explain some of the discrepancies between highly specific *in vitro* binding and broad binding highly overlapping targets *in vivo*. This metabolic role for IMP2 as well as its divergence from ZBP1 was clear from mouse studies, however high throughput studies often find significant overlap in the target pools. Reconciling the specificity differences of these proteins when comparing top down vs bottom up approaches remains a challenge and could benefit from orthogonal techniques to validate *in vivo* binding.

These studies^1,13,33^ of the IMP family as well as the data presented here show the necessity of the GG motif for all IMP family members. This provides an important consideration when performing next generation sequencing of IMP family targets. A number of CLIP studies have relied on RNAse T1, leading to cleavage of RNA after G nucleotides and G depleting the final sequencing dataset^37^. As the “GG” sequence is required for binding to targets, RNAses other than T1 should be used for studies of the IMP family to avoid depletion of the diguanosine motif in the final dataset. This may explain why in a previous study of the IMP family, only the CA motif was found to be enriched, instead of both the CA and GG motifs^14^. It would be possible to avoid this bias with newer methods that avoid the RNAse digestion and multiple processing steps of CLIP.

In addition, to avoiding sequencing bias, analysis of high throughput methods often fails to recognize the bipartite consensus element necessary for binding. The single consensus elements that have previously been discovered for the IMP family do not contain much complexity, often did not bind to canonical targets and did not explain the binding preferences of each protein. Given that the IMP family, like most RNA binding proteins rely on multiple RNA binding domains where each domain has its specific sequence preference, it is necessary for motif searches to consider possible randomized linker regions between specific binding motifs. Alongside previous work^13,33^, this study incorporates randomized variable linker regions between different domain consensus sequences when performing *in silico* target searches. The increased flexibility between consensus sequence elements helps discover targets of the IMP family and may inform future genome wide searches of other RBP targets.

In addition to discovering the consensus sequence of IMP2, this study validates the importance of the variable loop in determining specificity of closely related RNA KH domains. Studies have shown that the highly conserved GXXG loop is essential for binding to RNA^5,35^ and others have suggested that the variable region may be important for generating target specificity. There are several positively charged residues in the GXXG loops of the IMP family members and these regions may prefer to interact with the negatively charged phosphate backbone of the RNA, and from our structural data this would point to the variable loops functioning as readers of nucleotide identity. In addition to mutating the GXXG loops, studies have been able to mutate residues in the linker between KH domains^1^ or the KH domain variable loop^38^ to decrease the affinity of RBPs to their targets. This study provides the first gain of RNA-binding function mutations for the IMP family and for KH domains. While these variable loop mutations may be an IMP family specific it would be interesting to see if other families of KH domain containing proteins, such as NOVA, follow similar rules for generating specificity.

The SELEX experiments show that early on in selection, targets bound to both ZBP1KH34 and IMP2KH34 and contained a mix of nucleotides flanking the required ‘CA ‘and ‘GG’ motifs. Further selection with IMP2KH34 caused the SELEX pool to become more IMP2 specific. This correlated with enrichment of IMP2 specific sequences that dominated the pool (AGGU and UGGA), sequences that have been previously shown to prevent RNAs from binding to ZBP1KH34. This specificity was mediated solely by specific variable loop amino acid mutations.

It is likely that both the variable loops amino acids and target RNA sequences co-evolved when gene duplication of the IMP family occurred. Given that ZBP1 most resembles the ancestral family member and is essential for organism survival, the process of gene amplification of the IMP family members must strike a balance between loss of function and gain of function. During its evolution IMP2 has lost the ability to bind to the ß-actin zipcode and evolved away from a role in ß-actin localization to provide overall benefit to the organism by gain of function. Therefore, IMP2 has neo-functionalized with respect with ZBP1 to regulate transcripts involved in energy metabolism. Interestingly the IMP2KO mouse is resistant to diabetes through an increase in brown adipose tissue and increased thermogenesis. This phenotype is strikingly similar to the function of one of its evolutionarily conserved targets, GRB10 (Sup Fig. 7). As the IMPs are noted to be translational repressors, it is possible that GRB10 is upregulated in the knockouts. Future studies may dissect how a combination of IMP2KO and GRB10 upregulation may lead to anti-diabetic phenotypes.

When comparing our work with two other studies^1,33^ of the IMP family KH34 domains we see several evolutionary trends. Given that ZBP1 and IMP3 are more related to each other than IMP2 we find that they recognize a similar consensus sequence. Interestingly though, there appears to be a divergence where ZBP1 can recognize both the original and swapped orientations of its consensus but IMP2 and IMP3 bind to only one orientation which represents the swapped orientation of the zipcode consensus.

Like many other Type I KH domains, the overall structure of ZBP1KH34 and IMP2KH34 are very similar in the free state. As has been previously shown in the limited number structures containing of KH RNA complexes, the RNA bound state shows little change compared to the RNA free state^10,11^. Co-crystals are extremely useful to determine the residues and bases involved in binding. When looking at these structures, proteins similar to the IMP family (poly C RNA binding protein and NOVA2^10^) show extensive interactions of RNA with the variable loops. However, to date, the few didomain co-crystals do not show a single RNA spanning an interaction with both domains but rather the structures have one RNA hairpin per domain^39^. This is contrary to the biochemical evidence and points towards a limitation of crystallography.

The IMP family members are selectively expressed during different developmental time points. While ZBP1 and IMP3 are largely limited to embryonic expression, IMP2 is broadly expressed and its expression persists throughout life. Immortalized cell lines as well as cancers selectively overexpress different IMP family members and their overexpression correlates with growth rate^40^, invasive potential^41^ and patient prognosis^42^. Work has also shown that IMP overexpression alone is sufficient to increase invasive potential^43^ and that the IMP family are amongst the most up regulated RBPs across the TCGA^44^. What is still not understood is why these selective patterns of IMP expression provide benefit to the cells or tumors and the first step required to understand this is a faithful and unbiased profiling of RNA targets. The *in vitro* approach taken in this study is not limited by cell type and provides the framework necessary for follow up *in vivo* studies of the IMP family.

## Methods

### Plasmids and cloning

For purification of maltose binding protein fusions, the coding sequences for ZBP1KH34 (404–561) and IMP2KH34 were cloned into a modified pMalc2 where a C-terminal 6xHIS tag was added by PCR (Chao et al 2012).

For purification of his tagged protein fusions, the coding sequences for ZBP1KH34 (404–561) and IMP2KH34 were placed in a pet22HT vector, downstream of N terminal 6xHIS tag followed by a TEV site.

Mutations were introduced by synthesizing in frame DNA oligos containing the appropriate restriction sites (Invitrogen). Oligos were annealed by mixing in each oligo (1uM) in 100mM NaCl. The mixture was heated at 95°C for 5 minutes then allowed to cool on the bench top. This reaction was then diluted 100x and 1uL was used for ligations.

### Recombinant expression

Bl21 Rosetta 2 cells (EMD Millipore) were combined with the plasmid of interest and transformed by heat shock. Single colonies were selected for 5mL cultures and then transferred to 1L of either LB (for unlabeled protein expression) or M9 deuterated minimal media supplemented with 3g ^13^C, ^2^D glucose and 1g ^15^N NaCl (for NMR spectroscopy experiments) and incubated at 37°C. The cultures were monitored by OD600 readings and induced with 1mM IPTG when the OD600 reached between 0.6-0.8. Induction was performed for 4 hours at 37°C, at that time cells were pelleted and stored at −80°C until ready for purification.

MBP Protein preparation was performed by a crushed Complete EDTA-free protease inhibitor tablet (Roche) was added to the cell pellet before being resuspended in lysis buffer (50mM Tris pH 7.5, 1.5M NaCl, 1mM EDTA, 1mM DTT) and sonicated. Lysate was then centrifuged and the supernatant was passed over an amylose column (New England Biolabs) and washed for 4 hours. Fractions were eluted and checked by comassie stained SDS-Page gel. Protein was concentrated using centricon spin column, final protein concentration was determined by nanodrop absorbance at 280nm and/or by Bradford assay.

His Protein purification was performed by lysing bacterial cell pellets in 50mM sodium phosphate, 1.5M NaCl, 10mM imidazole and after centrifugation the supernatant was passed over a TALON column (Clontech). Fractions were eluted and checked by comassie stained SDS-Page gel. Protein was concentrated using centricon spin column, final protein concentration was determined by nanodrop absorbance at 280nm and/or by Bradford assay.

### EMSAs

100 pM RNAs were incubated at room temperature for 3 h with two-fold dilutions of the purified RNA binding protein in 10 mM Tris, 100 mM NaCl, 0.1 mM EDTA, 0.01 mg/mL tRNA, 50 μg/mL heparin and 0.01% IGEPAL CA630. Complexes were then run using 5% native PAGE in 0.5× TBE and visualized using the Typhoon 9400 variable mode laser scanner (GE Healthcare).

### NMR Spectroscopy, assignments and titration

NMR spectroscopy used purified His-IMP2KH34. 2D 1H 15N HSQC experiments were acquired on a Bruker DRX600MHz spectrometer. 3D deuterium decoupled gradient sensitivity enhanced triple resonance experiments [HNCO, HN(CA)CO, HNCA, HN(CO)CA, HN(CA)CB, and HN(COCA)CB] experiments were acquired either on a Varian INNOVA 600 or a Bruker Avance 800 spectrometer with non-uniform sampling. NMR data were processed using NMRpipe/NMRDraw^45^, analyzed using both CCPN Analysis^46^ and NMRFAM-Sparky^47^. Chemical shifts were indirectly referenced to sodium 2,2-dimethyl-2-silanepentane-5-sulfonate (DSS), using absolute frequency ratios for the 1H signal^48^.

### Crystallization, data collection and structure determination of IMP2 KH34

IMP2 KH34 (0.5mM) was crystallized using sitting-drop vapor diffusion at 22°C by mixing equal volumes of the protein and reservoir solution (23% PEG-3350, 200mM Ammonium sulfate, 100mM Tris pH 8.0). Crystals were cryoprotected by soaking them in the reservoir solution supplemented with 25% glycerol before flash cooling in liquid nitrogen. Data were collected to 2.05Å resolution from a single crystal at the Advanced Photon Source SGX-CAT beam line (Argonne, IL) at a wavelength of 0.9792Å. The diffraction data were indexed, integrated and scaled using Mosflm and the CCP4 suite of programs^49^.

The crystals of IMP2 KH34 belong to the monoclinic space group P2_1_ with unit cell parameters *a*=76.88, *b*=62.38, *c*=85.74, α=γ=90°, β=91.32°. The structure of IMP2 KH34 was determined by molecular replacement with Phaser using the structure of ZBP1 KH34 (3KRM) as a search morel^1,50^. The resulting model for IMP2 KH34 was then used as the input for the AutoBuild routine within Phenix for automated model building^51^. Rounds of manual model building and refinement were then performed using Coot^52^ and Phenix^53^. Protein stereochemistry was checked using Molprobity^54^. The final model contains residues 425-581 of IMP2 with 100% of amino acids in the allowed region and 98.58% of amino acids in the favored region of the Ramachandran plot.

### Systematic Evolution of Ligands by Exponential Enrichment (SELEX) procedure

Selection of RNAs was performed by preparing an antisense degenerate library with the sequence 5’-*TTTCGACGCACGCAACTATC*-(N30)-*GCTAAACTGCGTCGCTCTGCCC*-3’ by chemical synthesis (Integrated DNA Technologies). Thirty nucleotides (N30) were randomized to 25%A, 25%T, 25%G, and 25%C. Constant sequence regions are italicized. The N30 library (200 pmol) was converted to dsDNA by reverse transcription following the manufacturer’s specifications (SuperScript™ III Reverse Transcriptase; Invitrogen) using the IMP2N30 top strand primer 5’-GA<u>TAATACGACTCACTATAGG</u>GCAGAGCGACGCAGTTTAGC-3’. T7 promoter sequence is underlined. Then, N30 RNA pool library used for selection was generated by *in vitro* T7 transcription (MEGAshortscript™ T7 Transcription Kit; Ambion) of the dsDNA library and gel-purified. The stringency of selection was increased by progressively reducing the concentration of the purified MBP-IMP2KH34 in the binding reaction: in the first round of selection, MBP-IMP2KH34 (300 nM) was equilibrated with the library RNA pool (400 nM). In subsequent rounds of selection, the concentration of MBP-IMP2KH34 was reduced to 30 nM (rounds 2 and 3), 15 nM (round 4), 3 nM (round 5) and 1 nM (round 6-9) while RNA concentration was reduced to 300 nM (round 2-4), 90 nM (round 5), 30 nM (round 6) and 60 nM (round 7-9). After phenol/chloroform extraction, the RNA selected in every round was converted to cDNA by reverse transcription (SuperScript™ III Reverse Transcriptase; Invitrogen) using the IMP2N30 bottom strand primer 5’-TTTCGACGCACGCAACTATC-3’. The resulting cDNA library was then amplified by PCR using the IMP2N30 top strand and IMP2N30 bottom strand primers and then *in vitro* transcribed into RNA for the next round of selection. Subsequent rounds of selection were performed similarly, except that a negative selection step was included by incubating the RNA pool with amylose resin in the absence of protein to remove any RNAs with nonspecific affinity for the amylose resin. N30 Random RNA pool library and RNA from rounds 4, 7, 8 and 9 were fluorescein-labeled, and their affinity for MBP-IMP2KH34 and MBP-ZBP1KH34 (control) was quantified by EMSA.

### Sequencing of SELEX data

The N30 Random DNA pool and cDNA from rounds 7-to-9 were cloned into TOPO™ TA Cloning™ Kit (Invitrogen) following manufacturer’s specifications. Colonies from the transformation were then individually picked and plated for colony sequencing (Genewiz). At least 25 colonies were sequenced per round using M13R primer 5’-CAGGAAACAGCTATGAC-3’.

### Bioinformatics

FoldUTR3 tables were downloaded from the UCSC table browser for human (GRCh37/hg19) and mouse (NCBI36/MM8). UTRs were queried for the nucleotide sequence CUCAC-(N10-15)-(A/U)-GG-(A/U), where N represents any intervening nucleotide. Genes demonstrating conservation of this sequence motif between human and mouse genomes were identified as potential RNA ligands of IMP2. IPA Knowledge Base 9 (Ingenuity Systems) was used to determine diseases and functional category enrichment of these potential ligands. The P-value, based on a right-tailed Fisher’s exact test, considers the number of identified focus genes and the total number of molecules known to be associated with these categories in the IPA Knowledge Base. Significance was determined when an individual pathway reached a false discovery rate of <0.05.

Previously published eCLIP data for ZBP1 and IMP2 H9ES cells^15^ was visualized by retrieving input normalized tags and peaks for SEPT8 in the Integrated Genome Viewer.

RIP-seq was analyzed by downloading FASTQ files from HEK293T cells^36^ and mouse brown adipose tissue^17^ from SRA. FASTQ files were then aligned to a transcriptome index with kallisto^55^ and the output TPMs were used to perform Log2(RIP/control) calculations.

## Data Availability

Crystallography data (PDB-ID: 6ROL), NMR amide chemical shifts for KH3-4 (BMRB-ID: 27934) have been deposited. The source data underlying Figs. 2a, 2f and Sup Fig. 7 are provided as a Source Data file. All other relevant data are available from the authors upon request.

## Acknowledgements

The authors would like to thank members of the Singer lab for their helpful discussions and comments. They would also like to thank members of the Einstein NMR core facility (Mark Girvin and Sean Cahill) for help with NMR acquisition and for supervision of the data analysis. CE and RHS were supported by NIH grant R01NS083085. JB was supported with funding from the MSTP Training Grant T32GM007288 and predoctoral fellowship F30CA214009. NMR spectra were acquired on a Bruker 600MHz NMR instrument purchased using funds from NIH award 1S10OD016305 as well as a Bruker 80MHz NMR instrument at the New York Structural Biology Center.

## Author contributions

CE performed SELEX, analysis of Round 9 sequences and mutations of the IMP2 specific targets. VP performed genome wide search for IMP2 targets, JC performed X-ray crystallography of IMP2KH34; VB refined the crystal structure and modeled missing amino acids. JB designed and performed the experiments and subsequent analysis for all other figures. JB drafted the manuscript; JC, CE and RHS provided edits and feedback on the manuscript. CE and RHS supervised the research.

