## Supplementary Material for "The structural basis for RNA selectivity by the IMP family of RNA binding proteins"

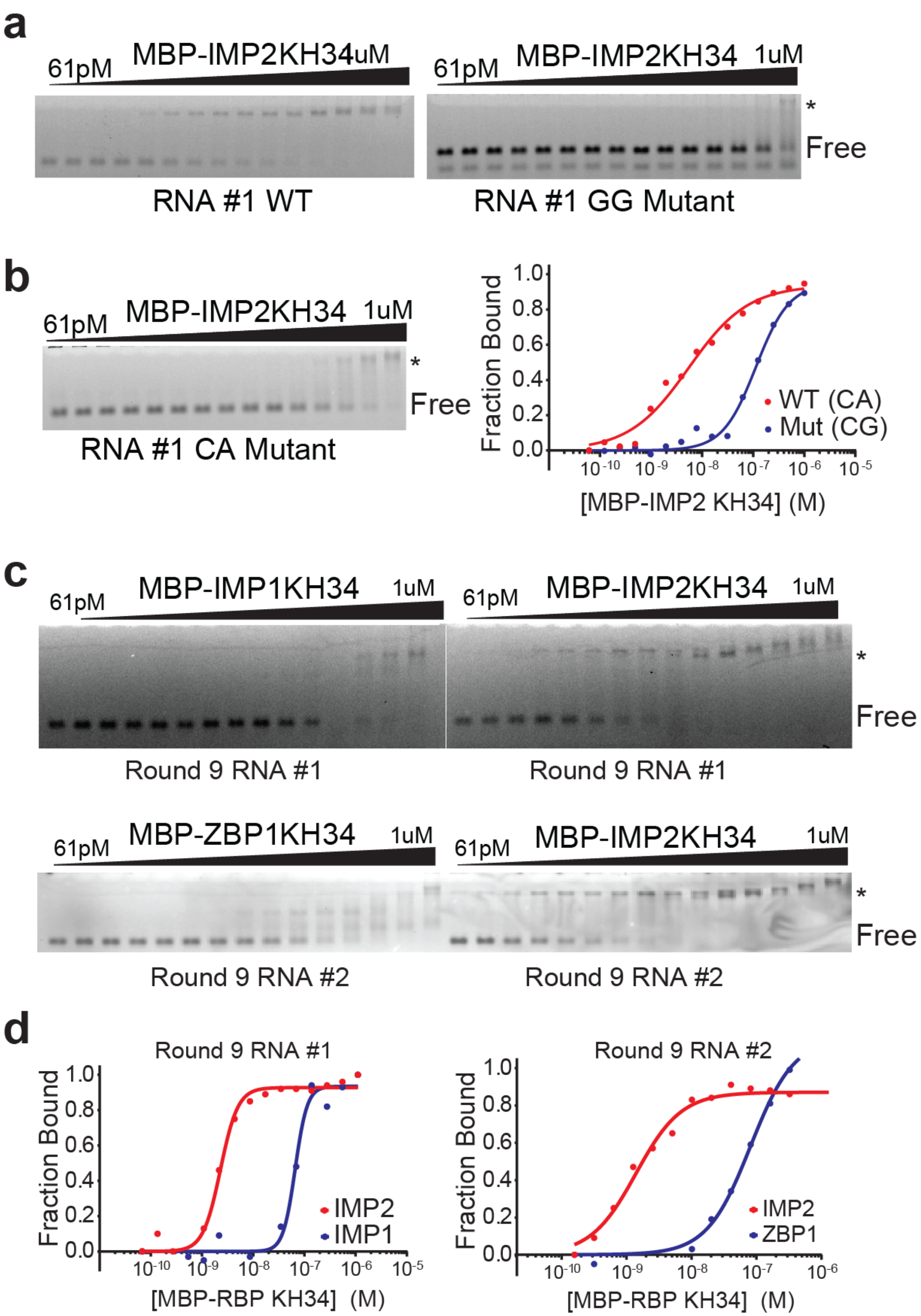
a
61pM
MBP-IMP2KH34uM
61pM
MBP-IMP2KH34
1uM
*
Free
RNA #1 WT
RNA #1 GG Mutant
b
61pM
MBP-IMP2KH34
1uM
*
Free
RNA #1 CA Mutant
Fraction Bound
WT (CA)
Mut (CG)
[MBP-IMP2 KH34] (M)
c
61pM
MBP-IMP1KH34
1uM
61pM
MBP-IMP2KH34
1uM
*
Free
Round 9 RNA #1
Round 9 RNA #1
61pM
MBP-ZBP1KH34
1uM
61pM
MBP-IMP2KH34
1uM
*
Free
Round 9 RNA #2
Round 9 RNA #2
d
Round 9 RNA #1
Fraction Bound
IMP2
IMP1
[MBP-RBP KH34] (M)
Round 9 RNA #2
Fraction Bound
IMP2
ZBP1
[MBP-RBP KH34] (M)

**Supplementary Fig. 1 Validation of IMP2 SELEX**

- **a.** Representative EMSA results for IMP2 KH34, ZBP1 KH34 and IMP2 KH34 binding to two RNAs from the round 9 SELEX pool. The filled triangle represents a 1:1 serial dilution of RBP KH34. The RBP KH34–RNA complex (*) and free RNA (FREE) are labeled.
- **b.** Quantification and fit to the Hill equation of EMSA results for IMP2KH34 (solid red line) and ZBP1KH34 (solid blue line) binding to the RNAs from panel A (two representative RNAs from the round 9 SELEX library pool).
- **c.** Representative EMSA results for IMP2 KH34 to WT RNA 1 and GG mutant RNA 1. The filled triangle represents a 1:1 serial dilution of IMP2 KH34. The IMP2 KH34–RNA complex (*) and free RNA (FREE) are labeled.
- **d.** Left - Representative EMSA results for IMP2 KH34 and CA mutant RNA #1. The filled triangle represents a 1:1 serial dilution of IMP2 KH34. The IMP2 KH34–RNA complex (*) and free RNA (FREE) are labeled. Right - Quantification and fit to the Hill equation of EMSA results for IMP2KH34 (solid red line) binding to WT (Red, data from Panel C) and CA mutant RNA #1 (Red).

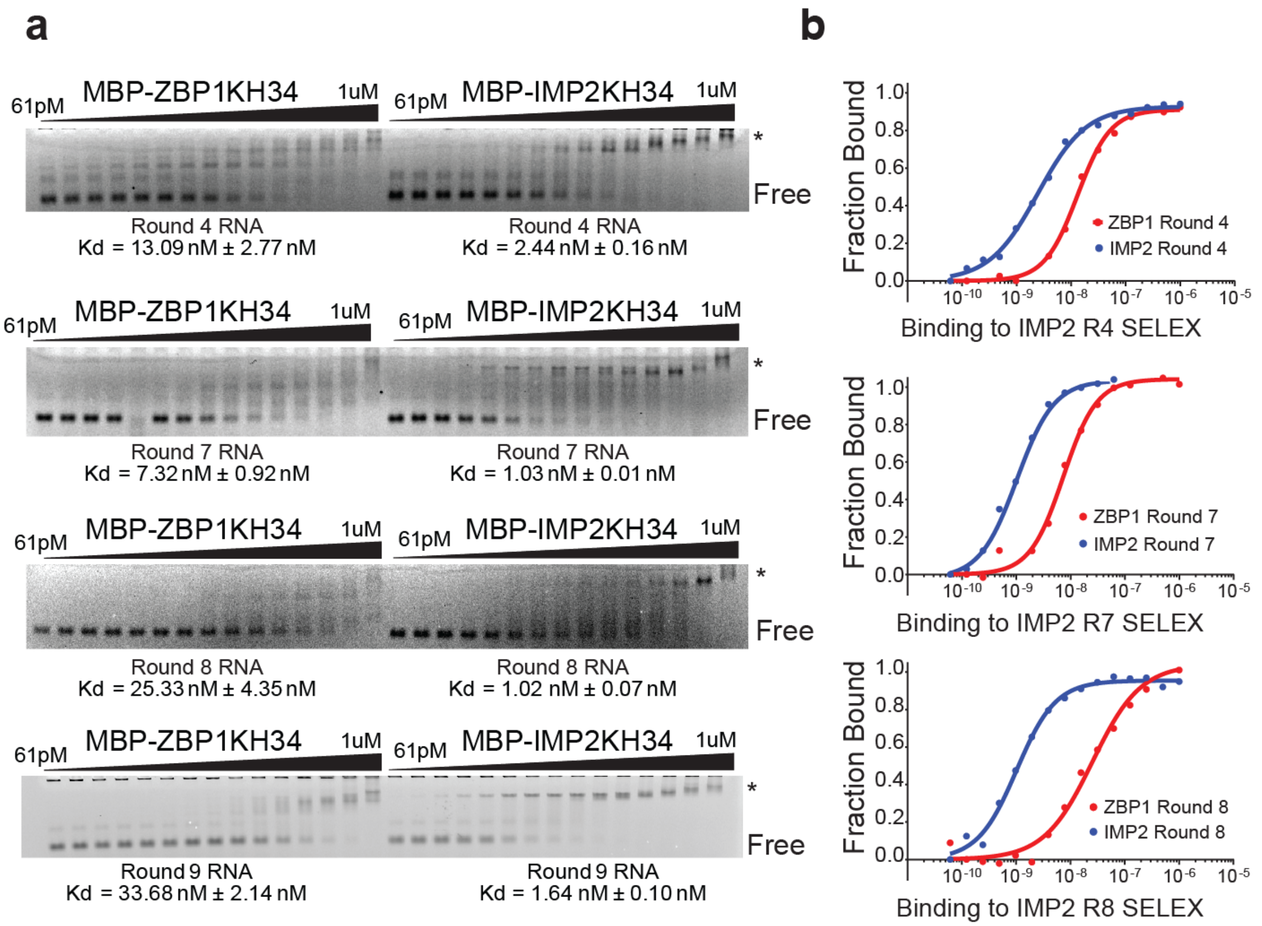


**Supplementary Fig. 2 IMP2 SELEX rounds tested against ZBP1 and IMP2**

- **a.** Representative EMSA results for ZBP1 KH34 (left) and IMP2 KH34 (right) binding to SELEX pools (rounds 4, 7,8 and 9 from top to bottom, respectively). IMP2KH34 Round 9 RNA gel from Fig 2B, used here for contrast with ZBP1KH34 Round 9 RNA. The filled triangle represents a 1:1 serial dilution of RBP KH34. The RBP KH34–RNA complex (*) and free RNA (FREE) are labeled.
- **b.** Quantification and fit to the Hill equation of EMSA results for IMP2KH34 (solid red line) and ZBP1KH34 (solid blue line) binding to the RNAs from panel A. The quantification of the round 9 SELEX EMSAs is found in Fig. 2C.

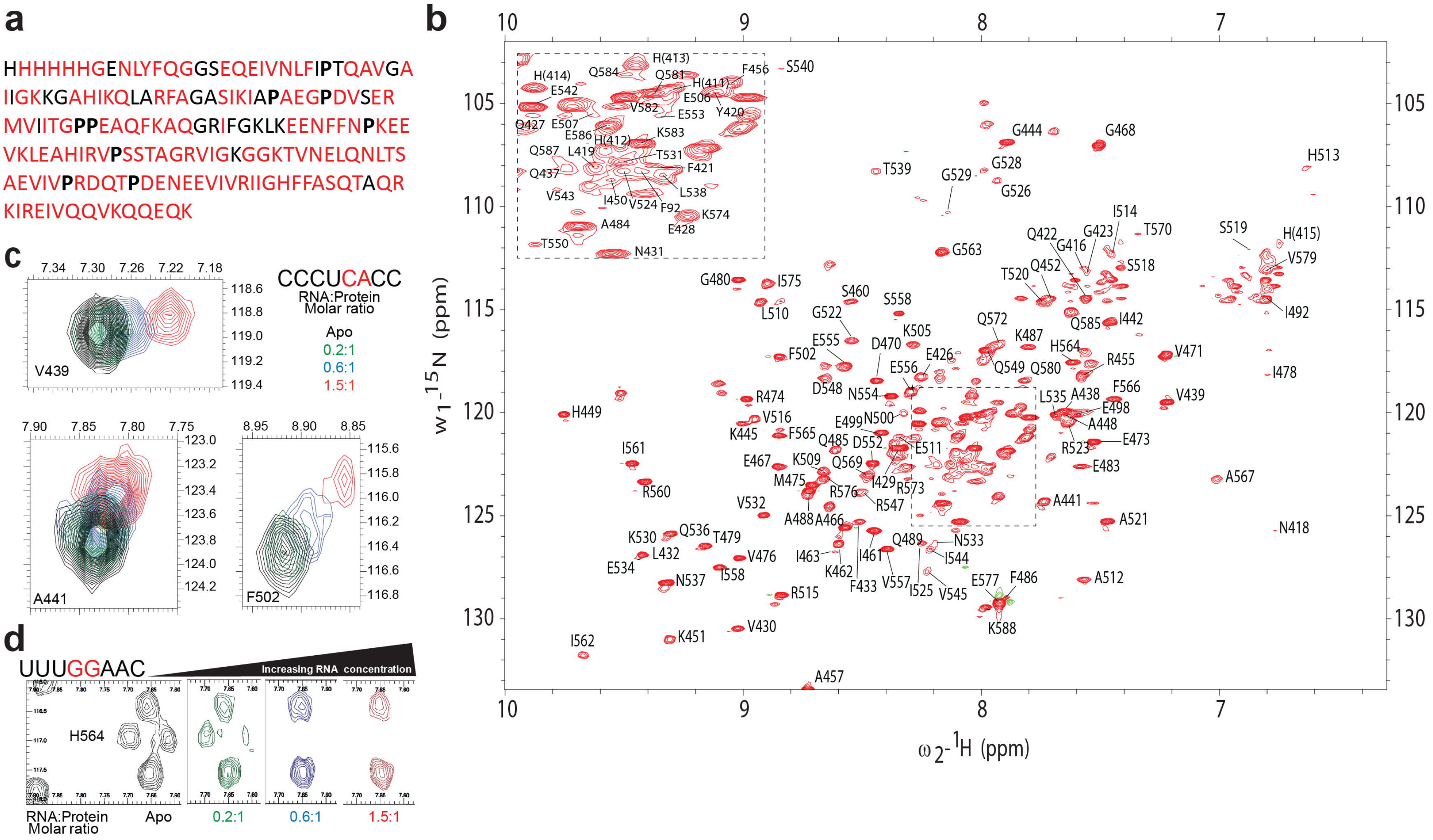
a
HHHHHHGENLYFQGGSEQEIVNLFIPTQAVGA
IIGKKGAHIKQLARFAGASIKIAPAEGPDVSER
MVIITGPPEAQFKAQGRIFGKLKEENFFNPKEE
VKLEAHIRVPSSTAGRVIGKGGKTVNELQNLTS
AEVIVPRDQTPDENEEVIVRIIGHFFASQTAQR
KIREIVQQVKQQEQK
b
w1-15N (ppm)
ω2-1H (ppm)
c
CCCUCACC
RNA:Protein Molar ratio
Apo
0.2:1
0.6:1
1.5:1
V439
A441
F502
d
UUUGGAAC
Increasing RNA concentration
H564
RNA:Protein Molar ratio
Apo
0.2:1
0.6:1
1.5:1

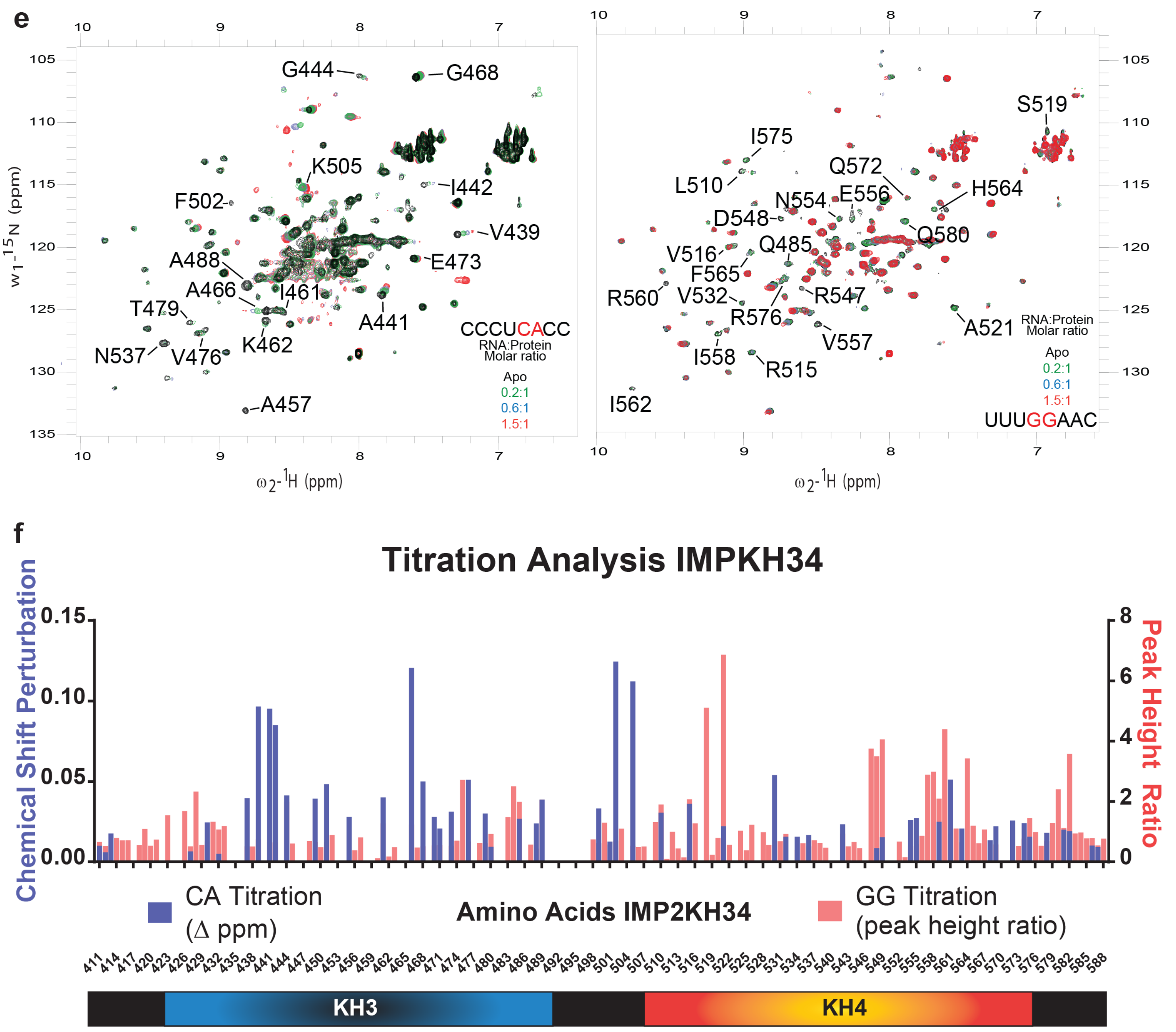
e
G444
G468
K505
I442
F502
V439
A488
E473
A466
I461
T479
A441
K462
N537
V476
A457
CCCUCACC
RNA:Protein
Molar ratio
Apo
0.2:1
0.6:1
1.5:1
S519
I575
Q572
L510
H564
N554
E556
D548
Q580
V516
Q485
F565
R560
V532
R547
R576
A521
V557
I558
R515
I562
UUUGGAAC
w1-15N (ppm)
ω2-1H (ppm)
f
Titration Analysis IMPKH34
Chemical Shift Perturbation
Peak Height Ratio
CA Titration
(Δ ppm)
Amino Acids IMP2KH34
GG Titration
(peak height ratio)
KH3
KH4

**Supplementary Fig. 3 Assigned spectrum of IMP2, examples of amino acid that are perturbed upon RNA titration**

- **a.** Successfully assigned amino acids in 6xHIS IMP2 KH34. Assigned residues are in red, unassigned are in black.
- **b.** Assignment of IMP2 KH34 amide-resonances. 15N-HSQC spectra of IMP2 KH34 labeled with peak assignments. Inset shows peak assignments within highlighted region (dashed box) of the spectra.
- **c.** Example of peaks during CA element titration, peaks are color coded according to the molar ratio of protein to RNA.
- **d.** Example of peaks during GG element titration, peaks are color coded according to the molar ratio of protein to RNA.
- **e.** 15N-HSQC spectra of IMP2 KH34 labeled with amino acids perturbed during RNA titrations, corresponding to Figure 3. RNA used for titration is noted on the bottom right of the spectra, with CA or GG element highlighted in red. Coloring of spectra corresponds to molar ratio of RNA to protein.
- **f.** Titration analysis of IMP2KH34 with CCCUCACC RNA (Blue, fast exchange, chemical shift perturbation on left axis) and UUUGGAAC RNA (Red, intermediate exchange, peak height ratio on right axis).

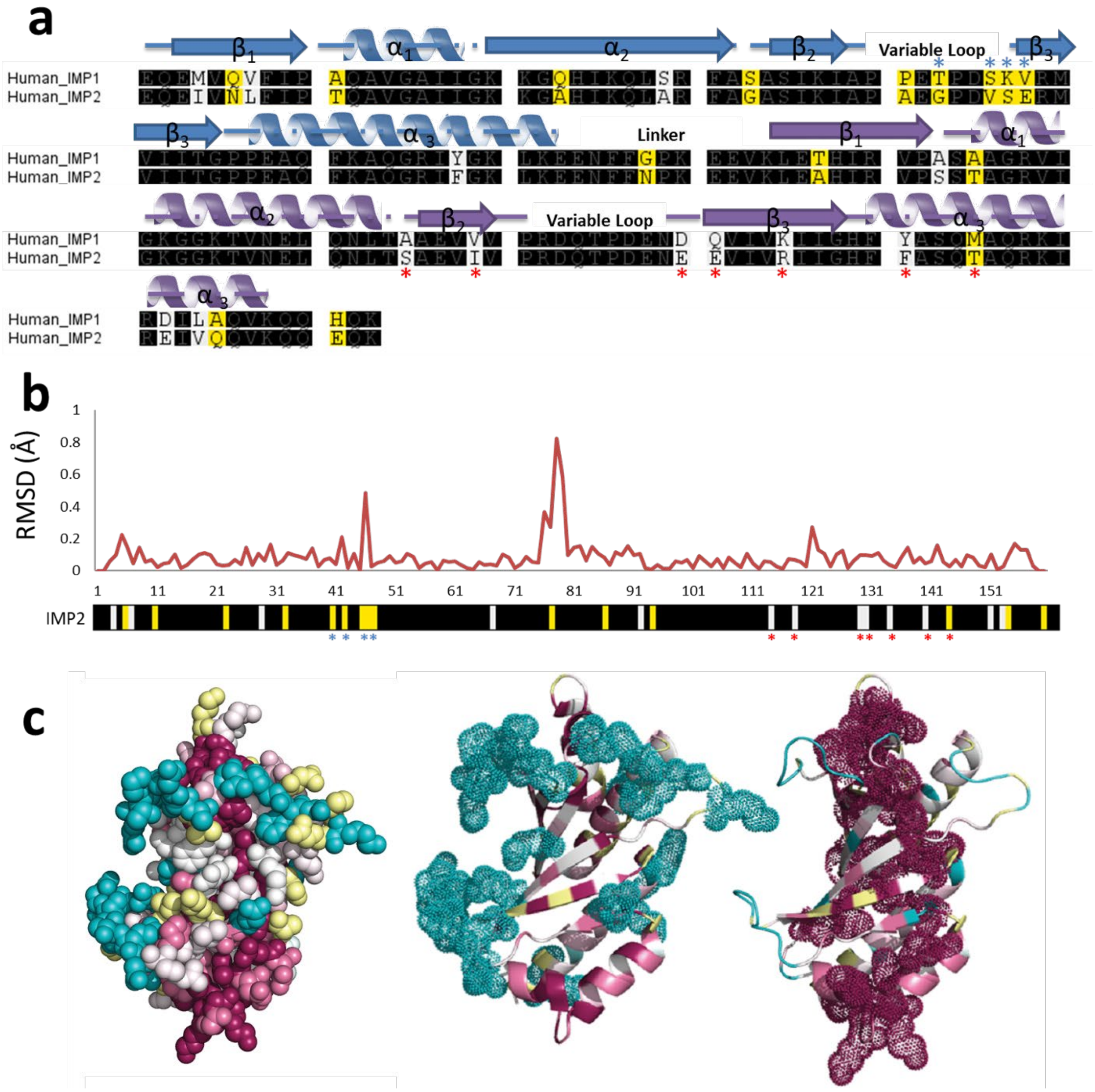


**Supplementary Fig. 4 amino acid and structural comparison of ZBP1 and IMP2**

a. Alpha helices and beta sheet topology of KH34 domains used to generate mutations. Black boxes represent conserved amino acids, yellow boxes represent non conserved amino acids with different chemical properties and white boxes represent non conserved amino acids with similar chemical properties. Mutations made in the KH3 (blue asterisks). Mutations made in KH4 (red asterisks).

b. RMSD differences between ZBP1 KH34 crystal structure (PDB: 3KRM) and IMP2 KH34 structure (this study) generated using AS2TS[59]. Below, sequence depiction of IMP2KH34 where black boxes represent conserved amino acids, yellow boxes represent non conserved amino acids with different chemical properties and white boxes represent non conserved amino acids with similar chemical properties. Mutations made in the KH3 variable loop (blue asterisks). Mutations made in KH4 variable loop (red asterisks).

c. Amino acid conservation across 50 IMP family member KH domains, generated with default ConSurf[60] parameters using PDBID 3KRM as a model. Amino acids are depicted as spheres and colored according to amino acid conservation, highly variable and highly conserved residues are in blue and red respectively.

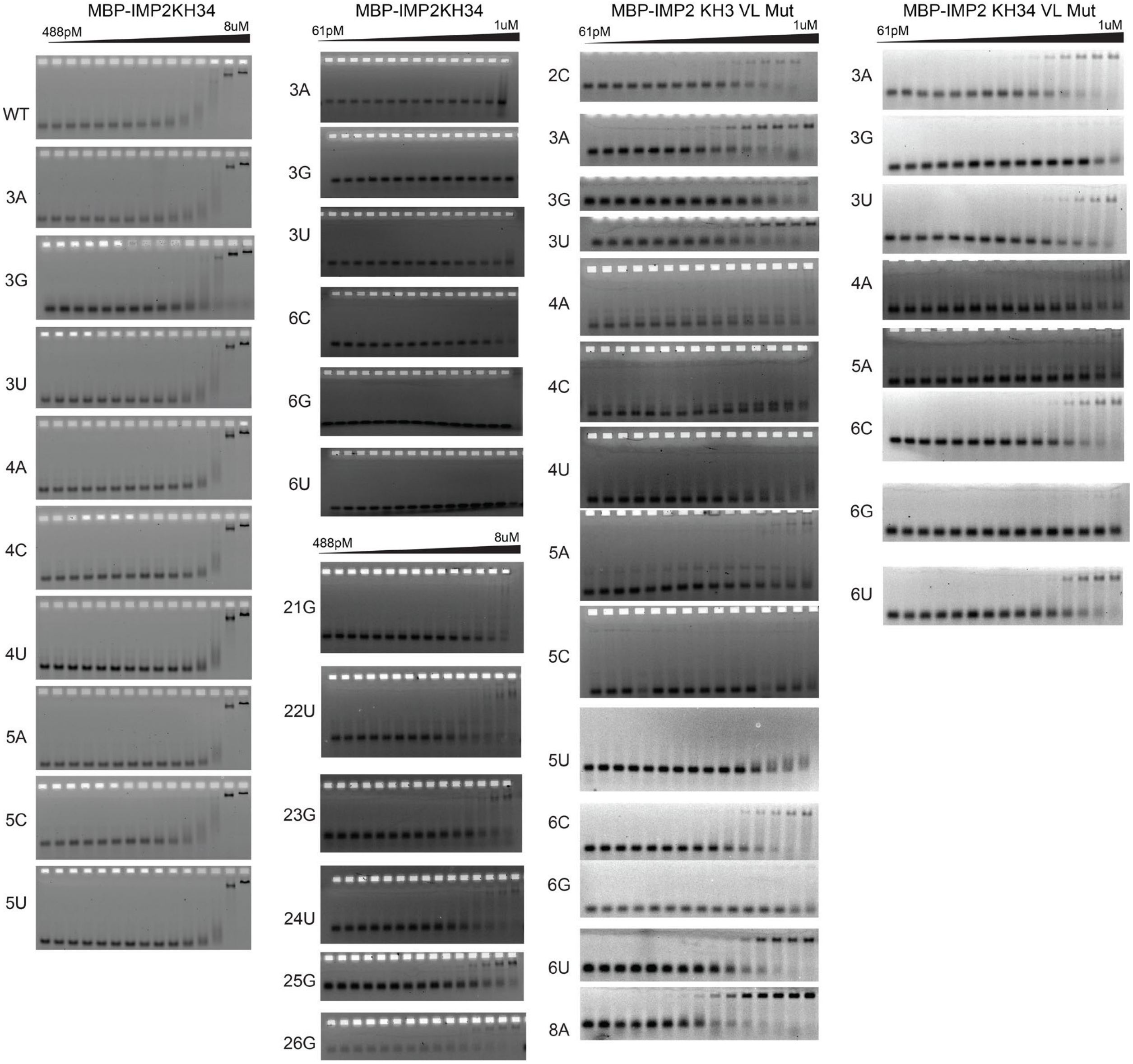


**Supplementary Fig. 5 Uncropped gels used in the manuscript**

a. Gels corresponding to Fig. 5A 5B and 5C.

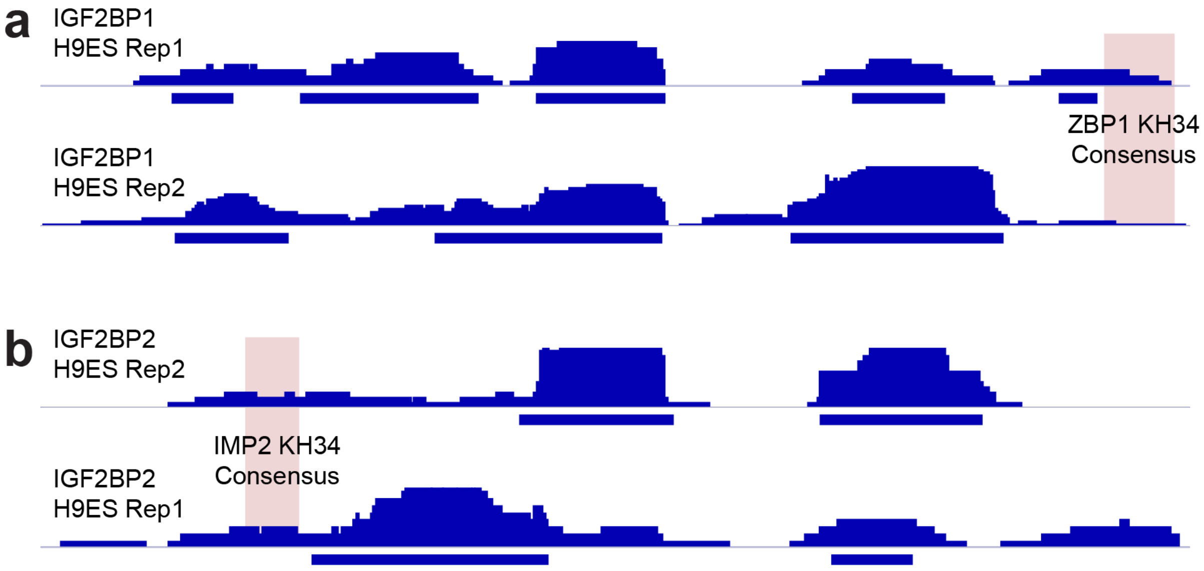


**Supplementary Fig. 6 Validation of consensus sequence targets with CLIP. SEPT8 is an RNA predicted to be co-regulated by ZBP1 and IMP2. Data from Conway et al., 2016.**

**a.** Above line - Replicate ZBP1 eCLIP tags for SEPT83' UTR. Below line – peaks called. Red box, consensus sequence for ZBP1KH34 (this study).

**b.** Above line - Replicate IMP2 eCLIP tags for SEPT83' UTR. Below line – peaks called. Red box, consensus sequence for IMP2KH34 (this study).

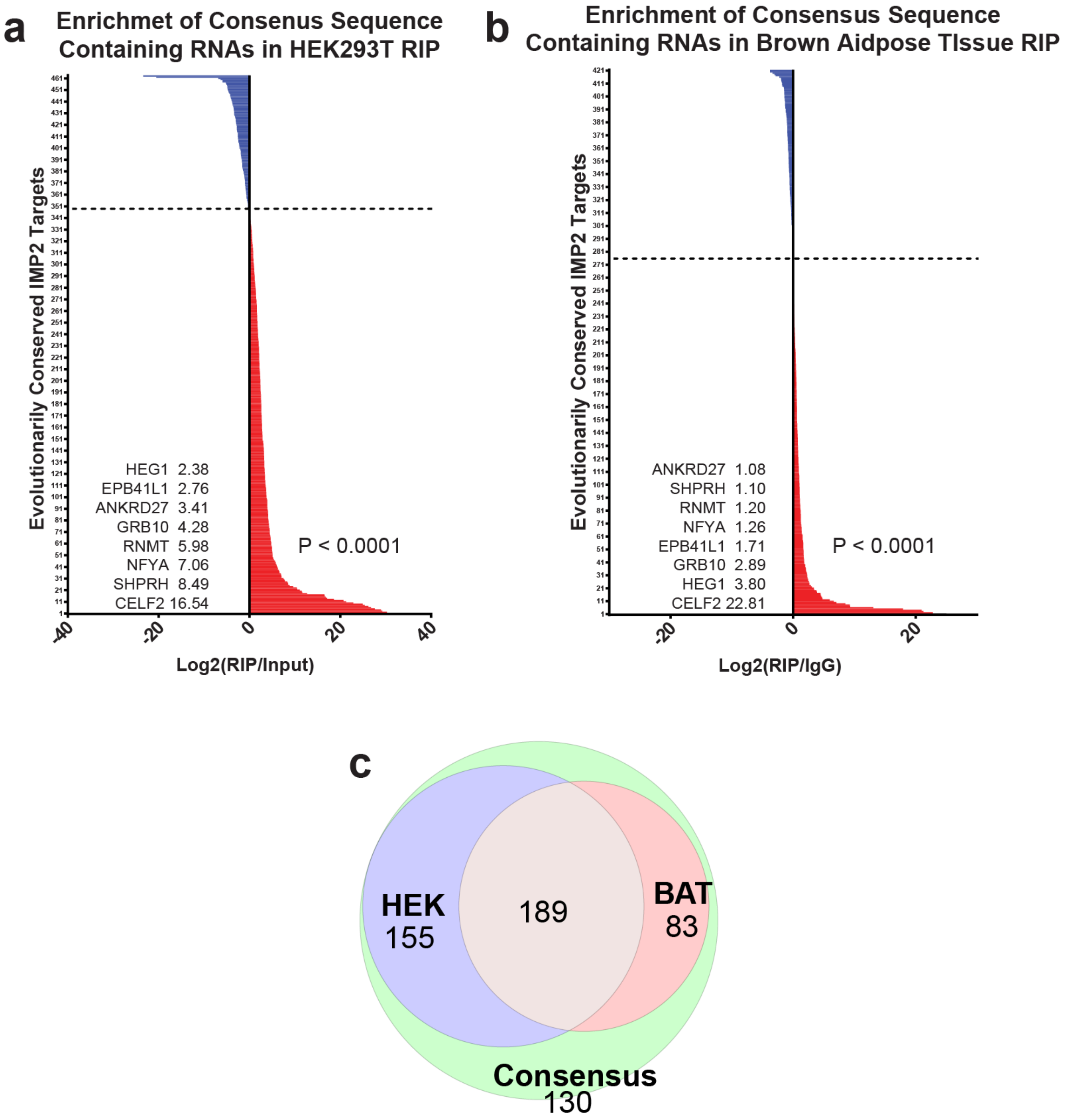


**Supplementary Fig. 7 Validation of conserved consensus sequence targets with RIP in human cells and mouse tissue.**

**a.** X axis is Log2(RIP/input) enrichment (red) or depletion (blue) of IMP2 RIP / Input after TPM normalization. Y axis are the rank ordered transcripts. Dashed line represents boundary between depletion or enrichment. Highly enriched genes with annotations related to diabetes listed with Log2 enrichment. P value was determined by single sample t test and shows significant enrichment of targets.

**b.** X axis is Log2(RIP/input) enrichment (red) or depletion (blue) of IMP2 RIP / IgG control after TPM normalization. Y axis are the rank ordered transcripts. Dashed line represents boundary between depletion or enrichment. Highly enriched genes with annotations related to diabetes listed with Log2 enrichment. P value was determined by single sample t test and shows significant enrichment of targets.

**c.** Venn diagram showing overlap of total high confidence consensus sequence containing RNAs (green circle), those found to be enriched in IMP2 RIP from brown adipose tissue (red circle) and those found to be enriched in IMP2 RIP from HEK293T cells (blue circle).

Source data are provided as a source data file.

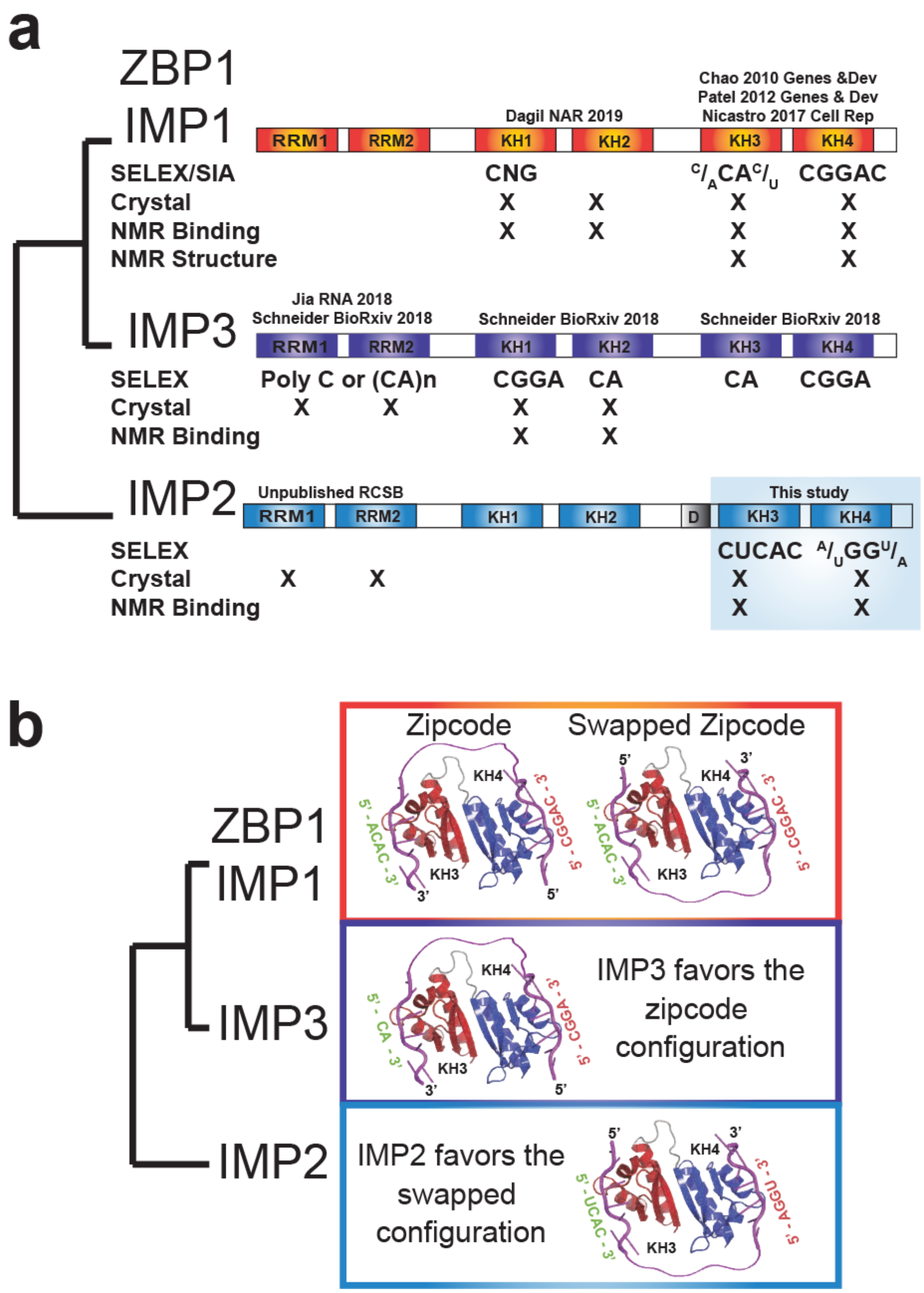


**Supplementary Fig. 8 Summary of IMP family structures, specificities and divergent evolution**

- **a.** Summary of structural and sequence studies performed and consensus sequences obtained to date on the IMP family.
- **b.** Summary of consensus orientations for the IMP family members.

| **Supplementary Table 1:** | IMP2 KH34 |
|---|---|
| **Data collection** | |
| Space group | $P2_1$ |
| Cell dimensions | |
| *a*, *b*, *c* (Å) | 76.88, 62.38, 85.74 |
| $\alpha, \beta, \gamma$ (°) | 90, 91.32, 90 |
| Resolution (Å) | 37.8-2.1 (2.175-2.1) |
| $R_{merge}$ | 0.0566 (0.4114) |
| $I / \sigma I$ | 8.05 (2.05) |
| Completeness (%) | 96.89 (98.74) |
| Redundancy | 1.9 (1.9) |

| | |
|---|---|
| **Refinement** | |
| Resolution (Å) | 37.8-2.1 |
| No. reflections | 46167 (4631) |
| $R_{work}$ / $R_{free}$ | 0.1981 / 0.2236 (0.2694 / 0.3026) |
| No. atoms | |
| Protein | 4907 |
| Ligand/ion | 198 |
| Water | 335 |
| *B*-factors | |
| Protein | 51.10 |
| Ligand/ion | 76.68 |
| Water | 51.51 |
| R.m.s. deviations | |
| Bond lengths (Å) | 0.005 |
| Bond angles (°) | 0.99 |
